# Convergent cortical temporal axis: common cortical oscillatory modes

**DOI:** 10.64898/2026.03.16.711992

**Authors:** Xiaobo Liu, Sanwang Wang, Xinyu Wu, Siyu Long, Liang He, Lang Liu, Ruifang Cui, Guoyuan Yang

**Affiliations:** School of Interdisciplinary Science, Beijing Institute of Technology, Beijing, China; McConnell Brain Imaging Centre, Montreal Neurological Institute, McGill University, Montreal, Canada; School of Medical Technology, Beijing Institute of Technology, Beijing, China; Department of Human Genetics, McGill University, Montreal, Canada; Peking University Sixth Hospital, Peking University Institute of Mental Health, NHC Key Laboratory of Mental Health (Peking University), National Clinical Research Center for Mental Disorders (Peking University Sixth Hospital), Beijing, 100191, China; School of Information Engineering, Huzhou University, No. 759, East Second Ring Road, Huzhou, 313000, Zhejiang, China

**Keywords:** neural oscillations, MEG, Spectral coordination modes, cortical hierarchy, excitation–inhibition balance, Multiscale brain organization, Parkinson’s disease

## Abstract

Brain dynamics provide the most direct substrate for neural function and representation. Extensive prior work has demonstrated that the human brain is organized into hierarchical structures across molecular, cellular, microstructural, and macroscale network levels. However, whether these hierarchies can be directly linked through shared patterns of neural dynamics remains unresolved, and clear evidence from data-driven analyses is still lacking. A central open question is whether brain dynamics vary along a single continuous dimension, or whether they are instead composed of a limited number of reproducible coordination modes that could provide a unifying basis for cross-scale integration.

Here, using source-resolved resting-state magnetoencephalography (MEG), we adopt a fully data-driven approach that imposes no a priori assumptions about cortical hierarchy or gradient structure. By characterizing full-spectrum power relationships across cortical regions, we identify a set of stable and reproducible spectral coordination modes. These modes capture how multiple frequency components jointly co-vary across space, defining distinct dynamical configurations rather than conventional band-limited oscillations or continuous gradients. The identified spectral coordination modes exhibit pronounced spatial differentiation across the cortex, distinguishing regions dominated by narrowband rhythmic activity from those characterized by broadband or multi-frequency coordination. Importantly, these modes do not correspond to predefined hierarchical axes, nor can they be reduced to existing MEG gradients or functional connectivity organizations. Instead, they emerge directly from the data, revealing a finite set of shared organizational patterns underlying cortical dynamics. Multimodal analyses further demonstrate that individual spectral coordination modes show selective associations with neurotransmitter system distributions, laminar microarchitecture, and cell-type–specific gene expression, suggesting that these dynamical patterns may serve as linking mechanisms between micro- and macroscale brain organization. Computational modeling supports this interpretation, showing that differences in local circuit parameters are sufficient to generate distinct spectral coordination states without invoking a single global dynamical hierarchy. Across the adult lifespan, these coordination modes exhibit frequency- and region-specific reorganization, indicating multiple parallel trajectories of dynamical aging. In Parkinson’s disease, alterations are observed in specific spectral coordination modes, particularly within higher-order cortical regions, suggesting that the disorder preferentially disrupts distinct dynamical configurations rather than inducing a global breakdown of cortical dynamics.

Together, these findings indicate that large-scale brain dynamics are not organized along a single pre-existing hierarchical axis, but instead are structured by a limited number of data-driven spectral coordination modes. This framework provides a new dynamical perspective for linking brain organization across modalities and scales, and offers a principled avenue for understanding systematic changes associated with aging and neurological disease.

## Introduction

From micro- to macro-scales, the human cerebral cortex exhibits a striking convergence of spatial organization: distributions of cell types and gene expression, microstructural features such as laminar differentiation and myelination, and macroscale phenotypes including functional connectivity and imaging-derived signatures all tend to align along a dominant organizational axis—the sensory–association axis (*1–8*). The two poles of this axis correspond, respectively, to sensory cortices specialized for input-driven, local processing and association cortices supporting transmodal integration and abstract representations (*8*). Thus, many seemingly heterogeneous cortical properties do not vary independently; instead, they co-vary in space around a common hierarchical principle.

If structure and connectivity provide the cortical “scaffold,” then brain dynamics constitute its most immediate, externalized expression in time (*9–11*). Oscillatory activity and related temporal processes directly reflect local circuit properties, long-range coupling, and the intrinsic timescales required for integration (*12*, *13*). Consequently, cortical dynamics should not be treated as fragmented frequency bands or purely local phenomena; rather, they are expected to exhibit cortex-wide, low-dimensional organization (*8*). This motivates a central question: if diverse cortical features converge on the sensory–association axis, do neural dynamics likewise converge onto a corresponding temporal axis—a systematic, low-dimensional progression of timescales and oscillatory features from sensory to association cortex?

Most work on macroscale axes has relied on fMRI or structural imaging, which are limited in their ability to directly characterize fast neural dynamics (*8*). In contrast, EEG/MEG provides millisecond-scale access to the spectral and rhythmic organization of cortical activity, offering a critical window into this putative temporal hierarchy (*14–16*). Importantly, the goal is not to define a “MEG gradient” as a modality-specific construct, but to identify a more general organizing principle: a convergent temporal axis of the brain—one that is stable across individuals, reproducible across datasets, and structurally aligned with the canonical sensory–association hierarchy.

Here, we use resting-state electromagnetic recordings to extract reproducible, cortex-wide common oscillatory patterns and organize them into an interpretable convergent temporal axis (and complementary dimensions). We then test whether this temporal axis aligns with established macroscale functional and imaging phenotypes and whether it can be anchored to microscale substrates, including cell-type and gene-expression topographies and laminar features. We aim to unify the cortex’s principal organizational axis in space with an overarching temporal axis of neural dynamics within a single cross-scale framework. Finally, we assess the reliability and clinical sensitivity of this temporal axis, evaluating its potential as a systems-level dynamical biomarker.

## Results

### Identification of cortex-wide common oscillatory modes

The primary exploration cohort was drawn from the Cam-CAN dataset and comprised 608 healthy adults (307 males, 301 females; mean age = 54.19 ± 18.19 years) who all completed resting-state MEG and MRI. We first constructed whole-cortex neurophysiological gradients from the MEG data. To this end, we (i) reconstructed source-level activity for each participant, (ii) estimated regional power spectra, and (iii) computed a spectrum-based similarity matrix using cosine similarity between cortical regions defined by the Schaefer 200-parcel atlas. Using source-reconstructed neural activity, we extracted cortical power spectra and applied the common orthogonal basis extraction (COBE) approach (*17*) to characterize dominant oscillatory features across regions. This procedure yielded frequency-resolved measures, including center frequency and center power, together with spatially coherent oscillatory patterns distributed across the cortex (Figure 1a).

**Figure 1.**
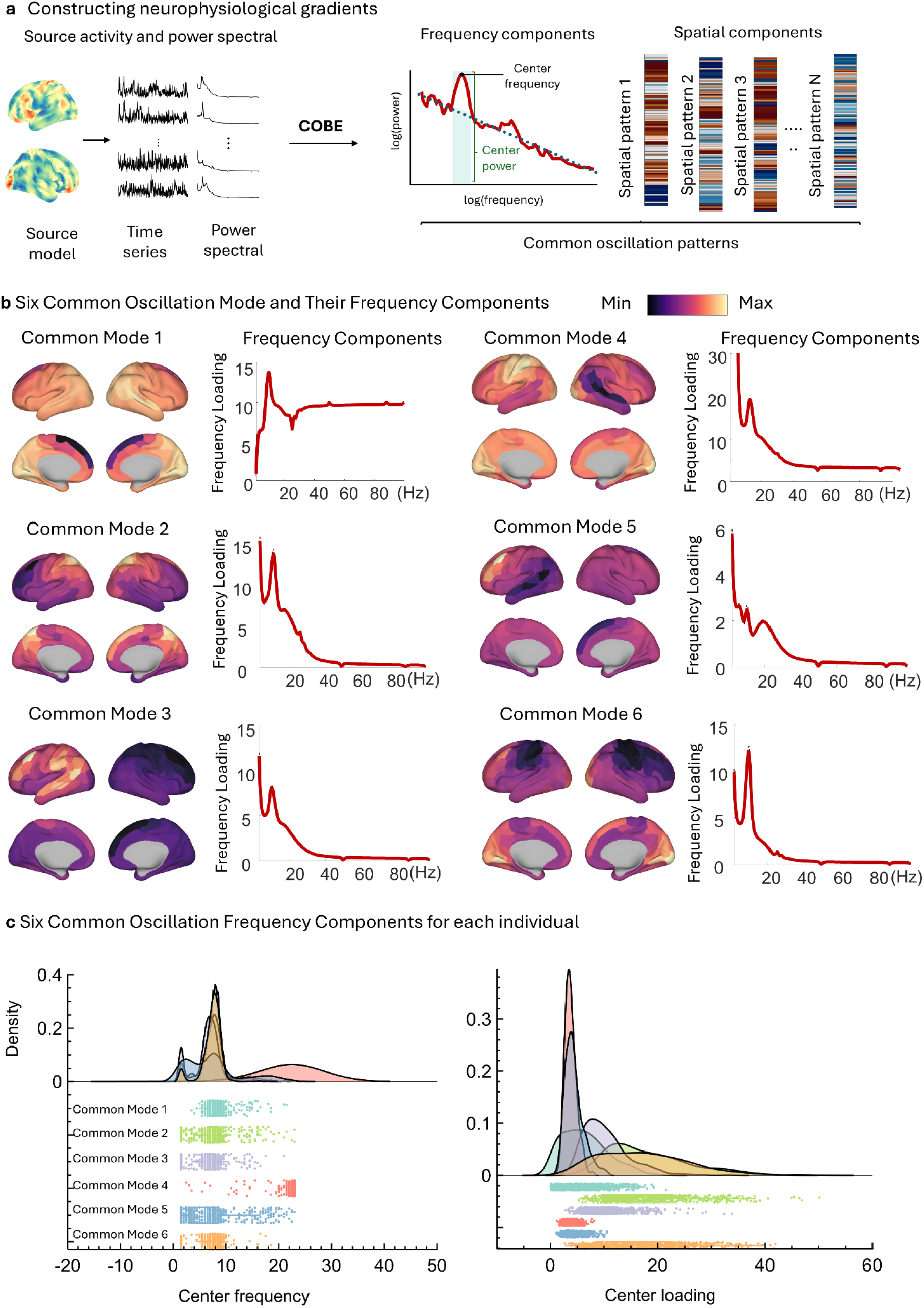
Construction of neurophysiological Patterns based on common oscillation patterns from -CAM-CAN dataset. **(a)** Schematic illustration of the pipeline for constructing neurophysiological gradients. Source-level neural activity was reconstructed and transformed into power spectra. The common orthogonal basis extraction (COBE) method was applied to extract frequency components, including center frequency and center power. Spatial components representing common oscillatory patterns across the cortex were subsequently identified. **(b)** Six common oscillation modes and their corresponding frequency components. For each mode, cortical surface maps illustrate the spatial distribution of oscillatory patterns, while line plots depict the associated frequency loadings across the spectrum. **(c)** Inter-individual distributions of center frequency (left) and center power (right) for the six common oscillation modes. Density plots and scatter distributions demonstrate variability across participants.

Decomposition of these features revealed six reproducible common oscillation modes at the group level (Figure 1b). Each mode exhibited a distinct spatial topography spanning multiple cortical systems and a characteristic frequency-loading profile, indicating separable axes of large-scale neurophysiological organization. These modes were consistently observed across participants, suggesting that they reflect fundamental properties of cortical dynamics rather than subject-specific noise.

Despite their robustness, the oscillation modes also showed marked inter-individual variability. Distributions of center frequency and center power demonstrated partially overlapping but distinguishable profiles across modes (Figure 1c), indicating that individual differences are embedded within a shared cortical oscillatory architecture.

### Alignment between common oscillation modes and multimodal cortical gradients

To determine whether oscillatory organization reflects established cortical hierarchies, we correlated common oscillation modes with previously described multimodal cortical gradients derived from functional connectivity, cortical geometry, and structure (Figure 2a).

**Figure 2.**
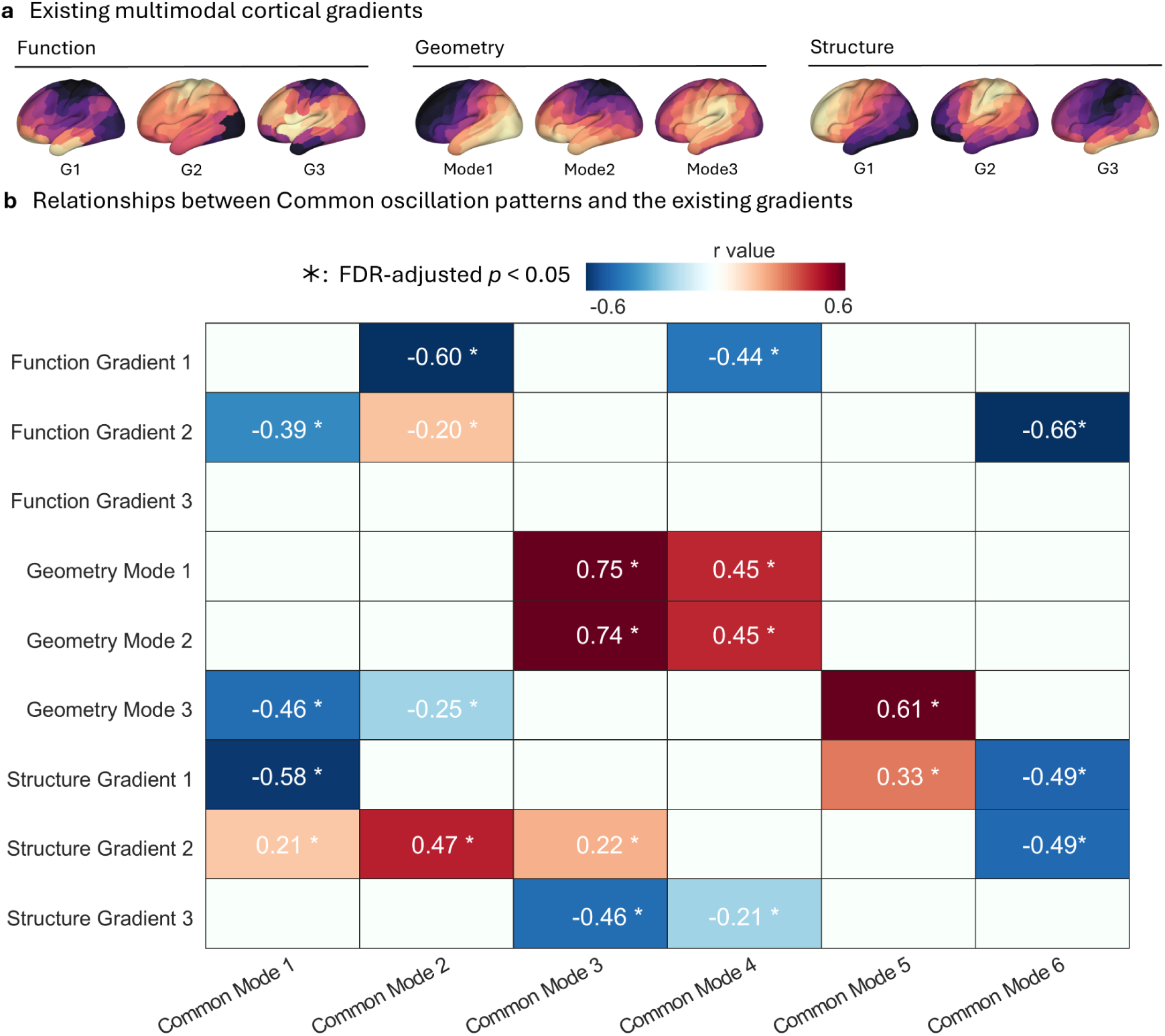
Relationships between common oscillation patterns and multimodal cortical gradients. **(a)** Previously established cortical gradients derived from functional connectivity (G1–G3), cortical geometry (Mode 1–Mode 3), and microstructural properties (G1–G3), displayed on the cortical surface. **(b)** Heatmap showing correlations between common oscillation modes and existing multimodal cortical gradients. Color indicates Pearson’s correlation coefficients (*r*), with asterisks denoting FDR-corrected significance (*p* < 0.05).

Significant associations were observed between oscillation modes and multiple gradients (Figure 2b). Specifically, **Function Gradient 1** was negatively correlated with Common Mode 2 (*r* = −0.60, FDR-adjusted *p* < 0.05) and Common Mode 4 (*r* = −0.44, FDR-adjusted *p* < 0.05). **Function Gradient 2** showed significant correlations with Common Mode 1 (*r* = −0.39, FDR-adjusted *p* < 0.05), Common Mode 2 (*r* = −0.20, FDR-adjusted *p* < 0.05), and Common Mode 6 (*r* = −0.66, FDR-adjusted *p* < 0.05).

Strong positive correlations were observed between geometric gradients and oscillation modes. **Geometry Mode 1** correlated with Common Mode 3 (*r* = 0.75, FDR-adjusted *p* < 0.05) and Common Mode 4 (*r* = 0.45, FDR-adjusted *p* < 0.05), while **Geometry Mode 2** showed similar associations (*r* = 0.74 and 0.45, respectively; FDR-adjusted *p* < 0.05). In contrast, **Geometry Mode 3** was negatively correlated with Common Mode 1 (*r* = −0.46, FDR-adjusted *p* < 0.05) and Common Mode 2 (*r* = −0.25, FDR-adjusted *p* < 0.05), but positively associated with Common Mode 5 (*r* = 0.61, FDR-adjusted *p* < 0.05).

Structural gradients also showed systematic relationships with oscillatory modes. **Structure Gradient 1** was negatively correlated with Common Mode 1 (*r* = −0.58, FDR-adjusted *p* < 0.05) and Common Mode 6 (*r* = −0.49, FDR-adjusted *p* < 0.05), whereas **Structure Gradient 2** exhibited positive associations with Common Mode 2 (*r* = 0.47, FDR-adjusted *p* < 0.05) and negative associations with Common Mode 6 (*r* = −0.49, FDR-adjusted *p* < 0.05).

### Associations between oscillatory gradients and cortical chemoarchitecture

We next examined the relationship between common oscillation modes and regional chemoarchitectural markers (Figure 3a). Multiple significant correlations were observed after FDR correction (*p* < 0.05).

**Figure 3.**
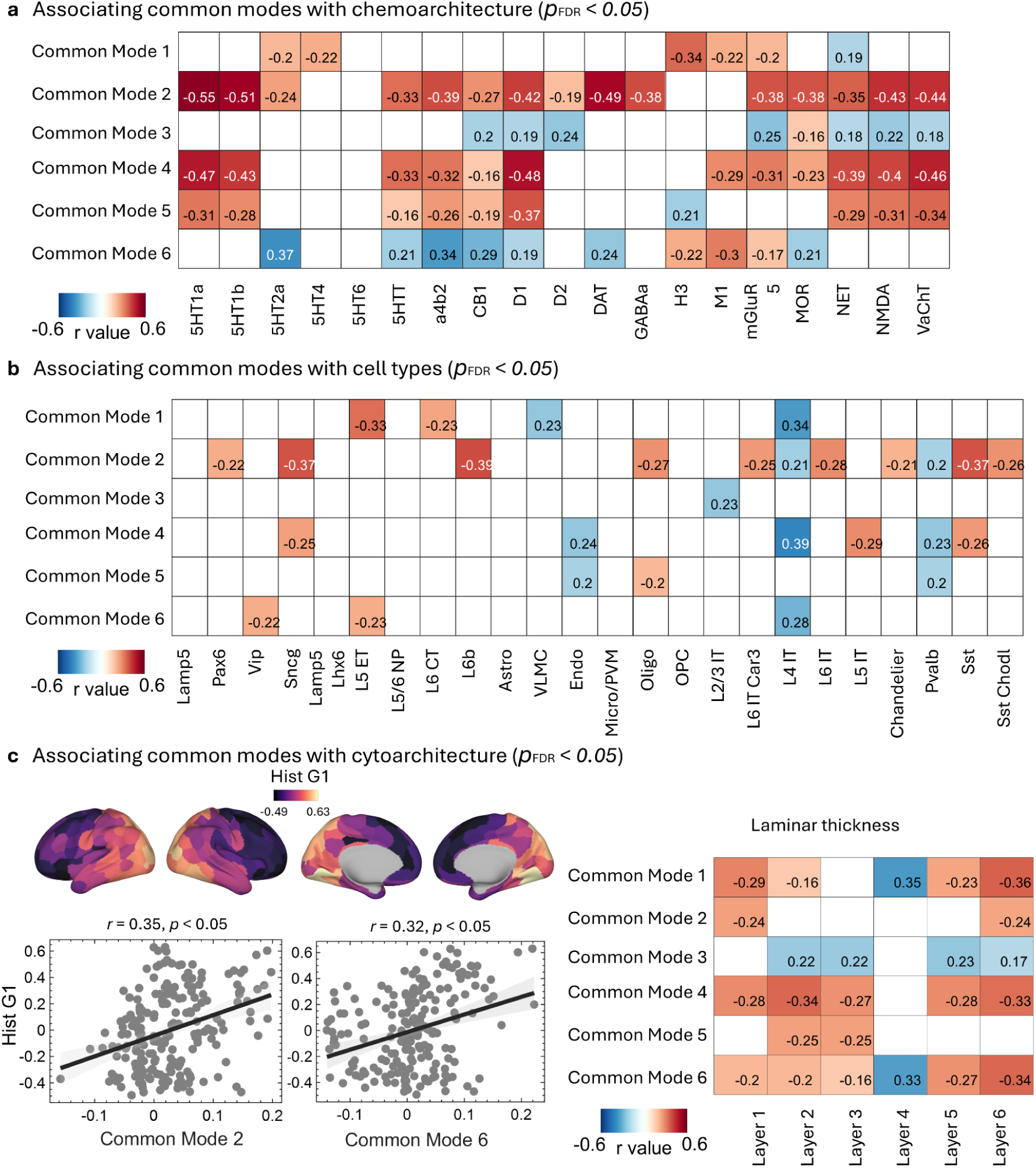
Associations between common oscillation modes and cortical microarchitecture. **(a)** Correlations between common oscillation modes and chemoarchitectural markers, including neurotransmitter receptor and transporter distributions. Only FDR-corrected significant associations (*p* < 0.05) are shown. **(b)** Associations between common oscillation modes and cortical cell-type distributions. Significant correlations (*p*<0.05, FDR-corrected) are displayed. **(c)** Relationships between common oscillation modes and cytoarchitectural features. Left: cortical maps and scatter plots illustrate associations with Histological Gradient 1 (Hist G1). Right: heatmap showing correlations between oscillation modes and laminar thickness across cortical layers.

Common Mode 2 showed widespread negative associations with serotonergic, dopaminergic, and GABAergic markers, including 5HT1A (*r* = −0.55), 5HT1B (*r* = −0.51), DAT (*r* = −0.49), and GABAA (*r* = −0.38). Common Mode 4 also demonstrated significant negative correlations with several neurotransmitter systems, including CB1 (*r* = −0.33), D2 (*r* = −0.48), and MOR (*r* = −0.39). In contrast, Common Mode 6 showed positive associations with selected dopaminergic markers (e.g., D1: *r* = 0.34; D2: *r* = 0.29).

### Associations with cortical cell-type distributions

Significant associations were also identified between oscillation modes and cortical cell-type distributions (Figure 3b). Common Mode 1 was negatively correlated with L5/6 IT neurons (*r* = −0.33, FDR-adjusted *p* < 0.05), whereas Common Mode 2 exhibited negative associations with Vip (*r* = −0.37) and L6 IT (*r* = −0.39).

Positive correlations were observed between Common Mode 4 and astrocytes (*r* = 0.24), as well as between Common Mode 5 and microglia/PVM (*r* = 0.20). Common Mode 6 showed positive associations with L4 IT (*r* = 0.28) and negative correlations with SST neurons (*r* = −0.26).

### Cytoarchitectural correlates of oscillatory gradients

Oscillation modes were further related to cytoarchitectural organization (Figure 3c). Histological Gradient 1 (Hist G1) was significantly correlated with Common Mode 2 (*r* = 0.35, *p* < 0.05) and Common Mode 6 (*r* = 0.32, *p* < 0.05).

Layer-specific analyses revealed significant correlations between oscillation modes and laminar thickness. For example, Common Mode 1 was negatively correlated with Layer 1 thickness (*r* = −0.29) and positively correlated with Layer 4 thickness (*r* = 0.35), whereas Common Mode 6 showed a strong positive association with Layer 4 (*r* = 0.33) and negative associations with Layers 5 and 6 (*r* = −0.27 and −0.34, respectively; all FDR-adjusted *p* < 0.05).

### Computational modeling links oscillatory gradients to excitation–inhibition balance

Using a Wilson–Cowan neural mass model constrained by the structural connectome, simulated center frequency maps were generated (Figure 4a). Simulated and empirical center frequencies were significantly correlated at the group level (*r* = 0.32, *p* < 0.05). At the individual level, the distribution of Spearman’s correlation coefficients was centered well above zero (median *r* ≈ 0.30), indicating consistent subject-specific correspondence.

**Figure 4.**
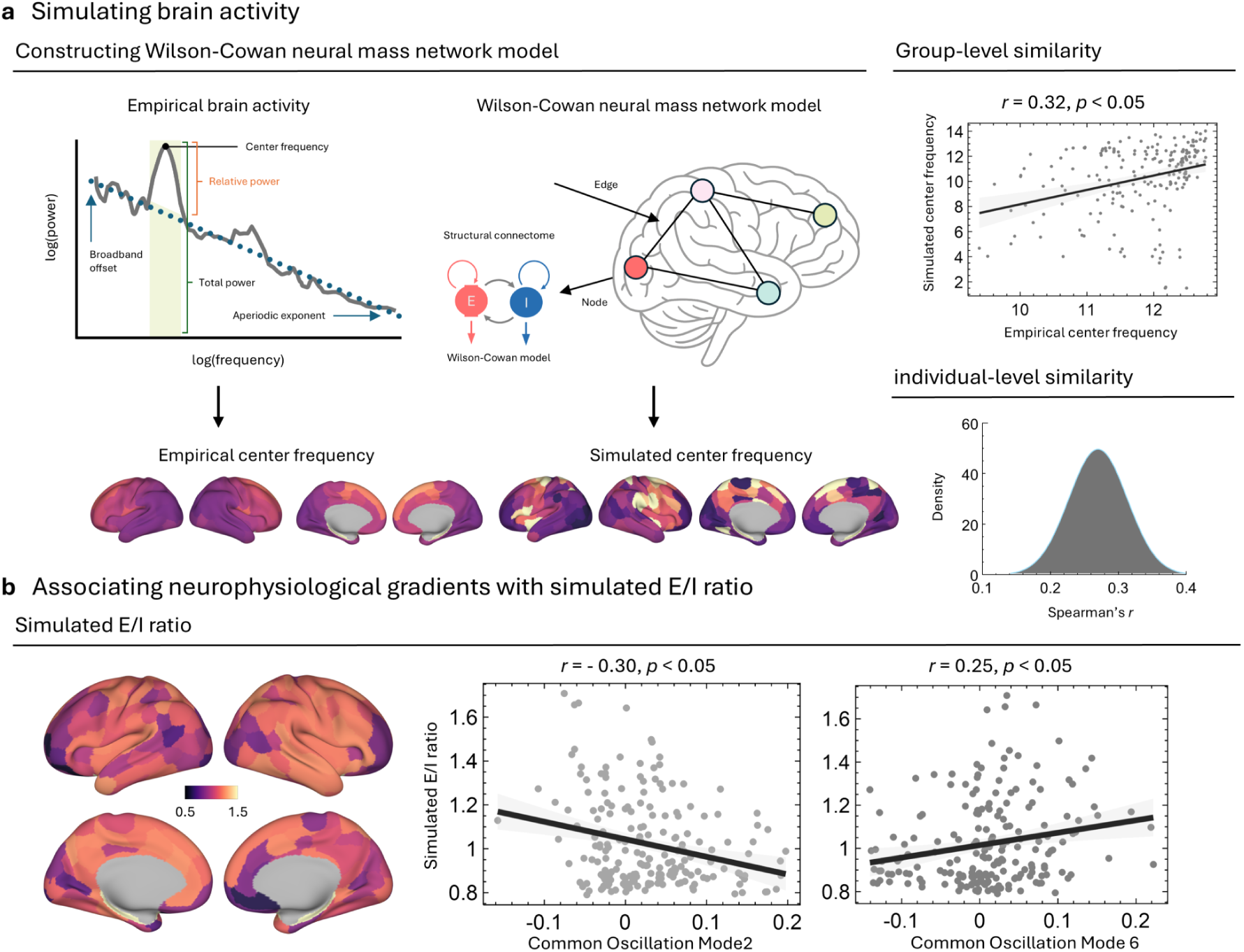
Linking neurophysiological gradients to computational modeling of brain activity. **(a)** Simulation framework based on the Wilson–Cowan neural mass model. Empirical power spectra were decomposed to extract center frequency, which was compared with simulated center frequency derived from the structural connectome. Group-level and individual-level similarities between empirical and simulated measures are shown. **(b)** Associations between neurophysiological gradients and simulated excitation–inhibition (E/I) ratio. Cortical maps display the spatial distribution of simulated E/I ratio, and scatter plots illustrate significant correlations with selected common oscillation modes.

Simulated excitation–inhibition (E/I) ratios showed significant associations with oscillation modes (Figure 4b). Common Mode 2 was negatively correlated with simulated E/I ratio (*r* = −0.30, *p* < 0.05), whereas Common Mode 6 showed a positive correlation (*r* = 0.25, *p* < 0.05).

### Altered oscillatory gradients in Parkinson’s disease

Group comparisons revealed significant alterations in oscillatory gradients in Parkinson’s disease relative to healthy controls (Figure 5). For Common Mode 2, patients exhibited significantly higher center frequency compared with controls (*t* = 3.54, *p* < 0.05). Common Mode 4 showed increased frequency loading strength (*t* = 2.44, *p* < 0.05), while Common Mode 6 demonstrated a significant shift in frequency distribution (*t* = 3.03, *p* < 0.05).

**Figure 5.**
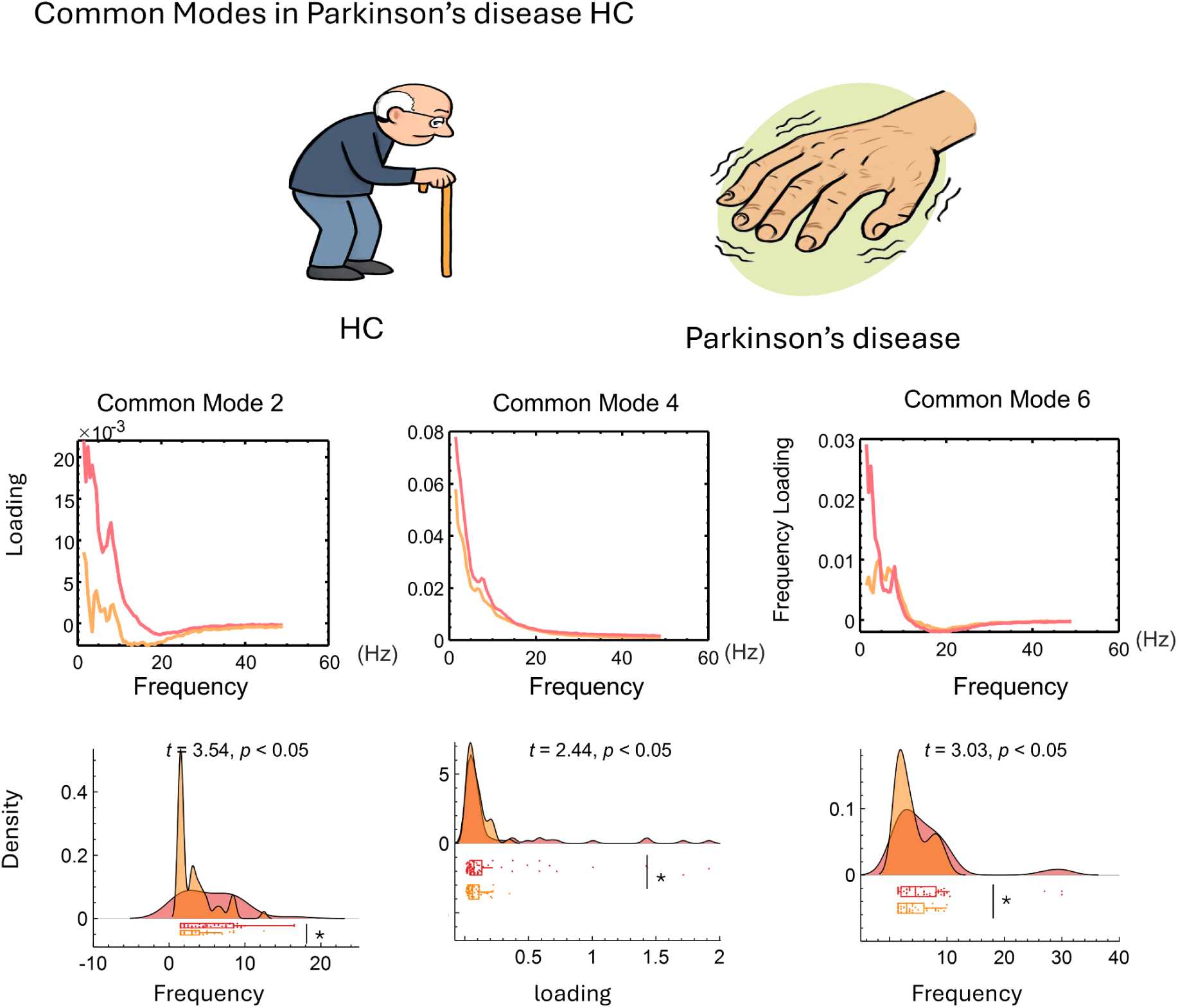
Alterations of common oscillation modes in Parkinson’s disease. Comparison of frequency characteristics of selected common oscillation modes between healthy controls (HC) and patients with Parkinson’s disease. Line plots show frequency loadings across the spectrum, while density plots illustrate group differences in center frequency or loading strength. Statistical significance was assessed using two-sample *t*-tests (*p* < 0.05).

#### Test–retest reliability of the frequency loadings for the six common oscillatory modes

Finally, we evaluated the reliability and external reproducibility of the MEG-derived gradients. Test–retest reliability was assessed using resting-state MEG data from the OMEGA dataset, including 158 unrelated participants (77 females; mean age = 31.9 ± 14.7 years), each contributing approximately 5 minutes of recording. We quantified the test–retest reliability of the frequency loadings associated with each of the six common oscillatory modes using intraclass correlation coefficients (ICCs). Across modes, ICC distributions were consistently skewed toward high values, indicating robust stability of the frequency-specific contributions across repeated measurements. Although some variability was observed between modes, the majority of frequencies exhibited moderate to high reliability. Collectively, these results demonstrate that the frequency loadings of the six common oscillatory modes exhibit high test–retest reliability, supporting their robustness as stable, individual-independent features of large-scale cortical dynamics.

**Fig. 6.**
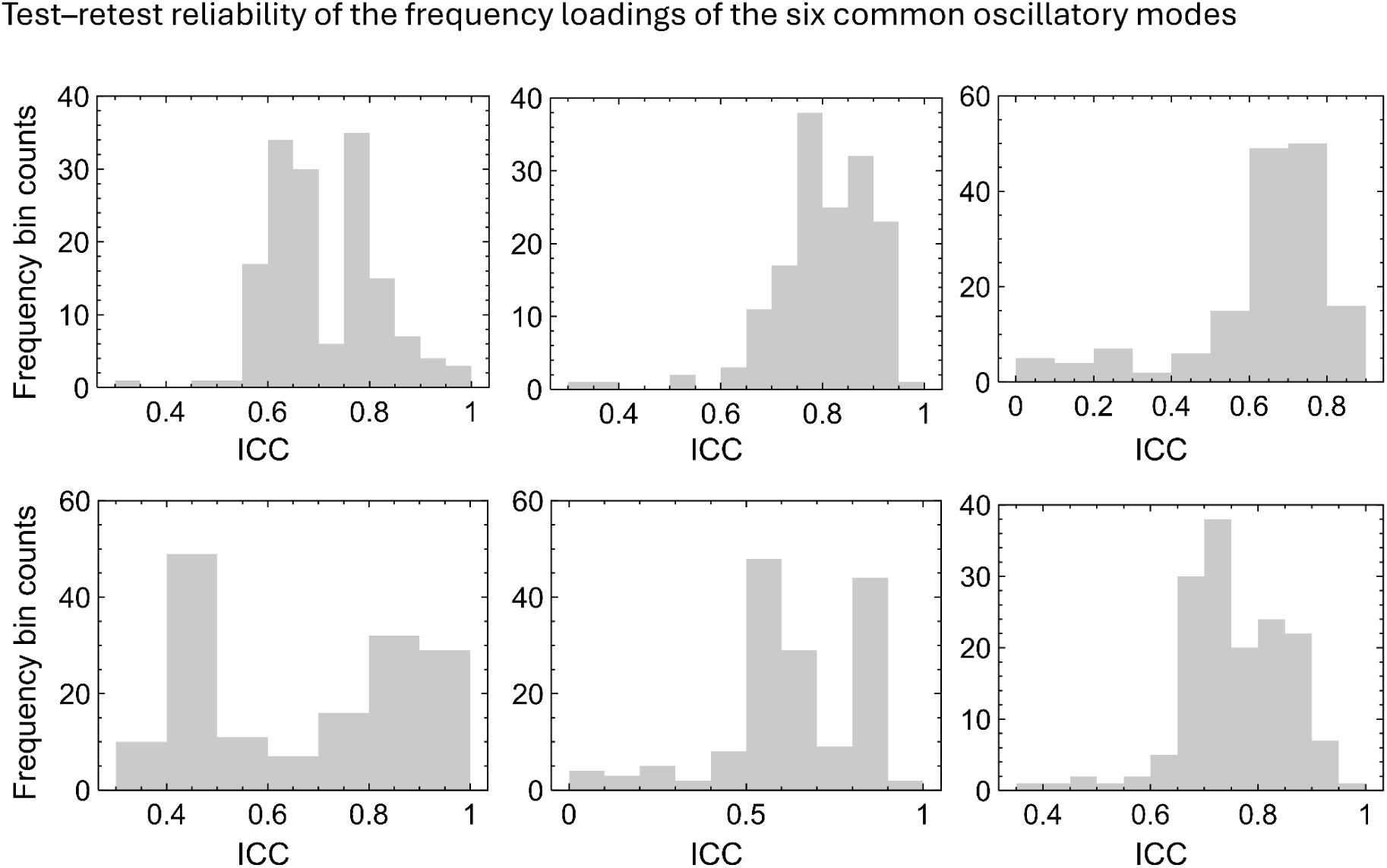
Test–retest reliability of the frequency loadings for the six common oscillatory modes. Each panel shows the distribution of intraclass correlation coefficients (ICC) across frequencies for one oscillatory mode. Histograms summarize the stability of frequency loadings across repeated measurements, with higher ICC values indicating greater test–retest reliability. Together, the results demonstrate that the frequency loadings of the common oscillatory modes are robust and reproducible across sessions, with variability in reliability across modes.

## Discussion

### Neurophysiological gradients as a unifying principle of cortical organization

In this study, we identified a set of six cortex-wide common oscillation modes that collectively form neurophysiological gradients across the human cortex. These gradients capture systematic variations in oscillatory dynamics that are spatially structured, reproducible across individuals, and aligned with established macroscale cortical hierarchies. By integrating electrophysiological features, multimodal cortical gradients, microarchitectural markers, and computational modeling, our findings suggest that large-scale oscillatory organization constitutes a fundamental dimension of cortical architecture.

Previous work has established functional connectivity, geometry, and microstructure as key organizing principles of the cortex. Our results extend this framework by demonstrating that oscillatory activity is not randomly distributed but follows structured gradients that recapitulate these canonical axes. Importantly, different oscillation modes selectively aligned with distinct functional, geometric, and structural gradients, indicating that cortical oscillatory organization is multidimensional rather than reducible to a single hierarchy.

### Linking oscillatory organization to cortical microarchitecture and cellular composition

A central finding of this study is the systematic association between neurophysiological gradients and cortical microarchitecture. Oscillation modes exhibited mode-specific relationships with neurotransmitter receptor and transporter distributions, as well as with cell-type and laminar features. These associations suggest that large-scale oscillatory patterns are constrained by regional variations in neurochemical signaling, cellular composition, and laminar differentiation.

The observed links to serotonergic, dopaminergic, and GABAergic systems are consistent with known roles of neuromodulators in shaping cortical excitability and rhythmic activity. Similarly, associations with excitatory and inhibitory neuronal populations, as well as glial cell markers, point to cellular substrates underlying macroscale oscillatory gradients. Together, these findings bridge microscale cortical architecture with macroscale neurophysiological dynamics.

### Excitation–inhibition balance as a mechanistic driver of oscillatory gradients

Using computational modeling, we demonstrated that empirically observed neurophysiological gradients can be partially reproduced by a Wilson–Cowan neural mass model constrained by the structural connectome. The correspondence between simulated and empirical center frequency patterns supports the notion that large-scale oscillatory organization emerges from interactions between network topology and local circuit dynamics.

Critically, different oscillation modes showed opposing relationships with the simulated excitation–inhibition (E/I) ratio. This finding suggests that regional differences in E/I balance may act as a key mechanistic driver of oscillatory gradients. Rather than reflecting uniform shifts in excitability, oscillatory organization appears to arise from structured variations in local circuit balance embedded within the connectome. This mechanistic interpretation provides a parsimonious explanation for the coexistence of multiple oscillatory gradients across the cortex.

### Disruption of neurophysiological gradients in Parkinson’s disease

Our results further demonstrate that neurophysiological gradients are altered in Parkinson’s disease. Patients exhibited significant changes in frequency characteristics of selected oscillation modes, indicating a disease-related reorganization of large-scale cortical dynamics. These alterations were particularly evident in modes dominated by low-frequency oscillations, consistent with previous reports of abnormal cortical and subcortical rhythmic activity in Parkinson’s disease.

Importantly, the observed changes extend beyond localized abnormalities and suggest a disruption of global neurophysiological organization. This gradient-level perspective provides a systems-level framework for understanding how Parkinson’s disease affects cortical dynamics and may help reconcile heterogeneous findings across electrophysiological studies.

### Conceptual implications and relation to prior work

Our findings build on and extend prior studies of cortical gradients by introducing neurophysiological gradients as a complementary organizational axis. Whereas previous work has focused on connectivity, microstructure, or gene expression, our approach integrates oscillatory dynamics into the gradient framework. This integration highlights oscillatory activity as a core feature of cortical organization, rather than a mere epiphenomenon of connectivity or structure.

Moreover, by linking oscillatory gradients to microarchitecture and E/I balance, our results provide a mechanistic bridge between electrophysiology and cortical anatomy. This framework may offer a unifying perspective for understanding how diverse neural processes—ranging from synaptic physiology to large-scale network dynamics—are coordinated across the cortex.

### Methodological considerations and limitations

Several limitations should be acknowledged. First, the oscillatory features were derived from source-reconstructed neural activity, which may be influenced by spatial leakage and model assumptions. Although we focused on robust, reproducible patterns, future work using invasive recordings could provide higher spatial specificity.

Second, associations with microarchitectural markers were based on group-level atlases rather than subject-specific measures. While this approach enables large-scale integration, individual variability in microstructure may further refine the observed relationships.

Third, the computational model employed a simplified representation of cortical dynamics and did not explicitly incorporate laminar or cell-type–specific mechanisms. More detailed models may better capture the full complexity of oscillatory gradients.

Finally, the Parkinson’s disease analysis was cross-sectional and focused on group differences. Longitudinal studies are needed to determine how neurophysiological gradients evolve with disease progression and treatment.

### Conclusions and future directions

In summary, we demonstrate that cortical oscillatory activity is organized along multiple neurophysiological gradients that align with canonical cortical hierarchies, reflect underlying microarchitecture, and are shaped by excitation–inhibition balance. These gradients are selectively disrupted in Parkinson’s disease, highlighting their clinical relevance.

Future work may extend this framework to other neurological and psychiatric disorders, investigate developmental and aging-related changes, and explore how neurophysiological gradients relate to cognition and behavior. By integrating oscillatory dynamics into the gradient framework, our study provides a comprehensive and mechanistically grounded view of large-scale brain organization.

## Method

### Exploration dataset

#### Cambridge Centre for Aging and Neuroscience Project Dataset

The exploration sample comprised 608 neurologically healthy adults (307 males, 301 females; mean age = 54.19 ± 18.19 years) drawn from the Cambridge Centre for Ageing and Neuroscience (Cam-CAN) cohort (*18*). All participants completed both resting-state magnetoencephalography (MEG) and structural and functional magnetic resonance imaging (MRI) acquisitions.

#### MEG Data Acquisition

Resting-state MEG data were collected at the MRC Cognition and Brain Sciences Unit (MRC-CBSU, Cambridge) using a 306-channel VectorView system (Elekta Neuromag, Helsinki), comprising 102 magnetometers and 204 orthogonal planar gradiometers housed in a magnetically shielded room (MSR). Signals were sampled at 1000 Hz and filtered online with a 0.03–330 Hz band-pass filter.

Head position within the MEG helmet was continuously monitored using four Head Position Indicator (HPI) coils, enabling offline correction of head motion. Vertical and horizontal electrooculogram (VEOG, HEOG) signals were recorded via bipolar electrodes to capture blinks and saccades, and an additional bipolar electrode pair recorded the electrocardiogram (ECG) to identify cardiac artifacts. The resting-state protocol yielded approximately 8 minutes and 40 seconds of continuous data per participant.

#### MRI Data Acquisition

Structural and functional MRI scans were acquired in a separate session at the same facility. High-resolution T1-weighted anatomical images and diffusion-weighted imaging (DWI) were obtained to support cortical surface reconstruction and atlas-based registration. Resting-state functional MRI (rs-fMRI) was collected using a gradient-echo echo-planar imaging (EPI) sequence with the following parameters: repetition time (TR) = 1970 ms, echo time (TE) = 30 ms, flip angle = 78°, field of view (FOV) = 192 × 192 mm², voxel size = 3 × 3 × 4.44 mm³, and 32 axial slices covering the cerebellum. Each rs-fMRI run lasted approximately 8 minutes and 40 seconds.

An additional multi-echo EPI sequence was acquired to enhance signal-to-noise ratio (TR = 2470 ms; TEs = 9.4, 21.2, 33, 45, and 57 ms), using the same spatial resolution and slice coverage. To correct for susceptibility-induced geometric distortions, dual-echo gradient-echo field maps were acquired using a spoiled gradient recalled echo (SPGR) sequence (TE1 = 5.19 ms, TE2 = 7.65 ms), with a total acquisition time of 54 seconds.

### Replication dataset

#### The Open MEG Archives

The replication sample was drawn from the Open MEG Archives (OMEGA) (*19*) and consisted of resting-state MEG recordings acquired on a uniform MEG platform (275-channel whole-head CTF system; Port Coquitlam, British Columbia, Canada). Data were sampled at 2400 Hz with a low-pass anti-aliasing filter at 600 Hz, and a built-in third-order spatial gradient noise cancellation scheme was applied.

Resting-state MEG data from 158 unrelated participants (77 females; mean age = 31.9 ± 14.7 years) were included. Each session comprised approximately 5 minutes of continuous recording. The participants underwent multiple sessions on separate days and contributed to the longitudinal fingerprinting analyses. All procedures were approved by the Research Ethics Board of the Montreal Neurological Institute, and all research activities adhered to the ethical guidelines of the institution.

### Parkinson’s disease dataset

#### The Open MEG Archives

This study analyzed resting-state MEG data from patients with Parkinson’s disease (PD) obtained from the OMEGA dataset, all of which were acquired using a uniform MEG system identical to that employed in the replication dataset.

The sample comprised healthy control participants (n=50n = 50n=50; mean age =60.58±6.83= 60.58 \pm 6.83=60.58±6.83 years) and individuals with idiopathic PD (n=65n = 65n=65; mean age =64.68±7.93= 64.68 \pm 7.93=64.68±7.93 years) at mild to moderate disease stages, as defined by Hoehn and Yahr scores ranging from 1 to 3. Participants were recruited through the Quebec Parkinson Network and underwent extensive clinical, neuroimaging, neuropsychological, and biological assessments. All patients with PD were receiving a stable antiparkinsonian medication regimen and demonstrated satisfactory clinical response prior to enrollment in the study.

#### MEG Data Preprocessing

Preprocessing and analysis of the MEG data were carried out using *Brainstorm* (2016 version) (*20*), employing default settings unless otherwise specified and in accordance with established best-practice guidelines. Initial preprocessing involved removal of slow drifts and DC offsets with a high-pass finite impulse response (FIR) filter at 0.3 Hz. Raw MEG traces were visually inspected to identify and exclude bad channels and time segments contaminated by prominent artifacts. Power-line interference at 50 Hz and its harmonics was attenuated using notch filters. A subsequent low-pass filter (0.6–280 Hz) was applied to suppress high-frequency noise while preserving physiologically relevant activity.

Physiological artifacts, including cardiac and ocular components, were removed using signal-space projection (SSP). Artifact-related components were identified with reference to ECG and EOG recordings. The cleaned continuous data were then segmented into non-overlapping 30-second epochs to enhance the robustness of resting-state analyses. These segments were band-pass filtered to the frequency range of interest and resampled at four times the upper cutoff frequency to minimize aliasing while maintaining signal fidelity.

Source reconstruction was performed using each participant’s T1-weighted anatomical MRI acquired on a 1.5T Siemens Sonata scanner. Cortical surface reconstruction and tissue segmentation were conducted using *FreeSurfer* (*21*). Cortical surfaces were tessellated into triangular meshes and visualized on inflated representations. Co-registration between MEG sensor space and individual anatomy was achieved using digitized scalp landmarks obtained during the MEG session. Forward modeling was implemented using the overlapping spheres method, and cortical source activity was estimated via linearly constrained minimum variance (LCMV) beamforming. Noise covariance matrices derived from empty-room recordings were used for normalization, thereby mitigating depth bias in source estimates.

#### Common Oscillation Modes Construction

Whole-cortex power spectra were computed at the vertex level and subsequently parcellated according to the Schaefer 200-region cortical atlas using CAM-CAN dataset. Principal component analysis (PCA) was then applied within each region to extract regional spectral sequences, thereby reducing spectral dimensionality while preserving dominant oscillatory features. In addition, the Common Orthogonal Basis Extraction (COBE) method (*17*) was applied to whole-brain spectral sequences across participants.

COBE identifies a set of orthogonal bases shared across individuals, enabling efficient low-dimensional representations of whole-brain spectral sequences and revealing cortical common oscillatory modes consistently expressed at the group level. At the same time, COBE separates individual-specific spectral variability into complementary subspaces, allowing stable group-level oscillatory structures to be characterized while preserving inter-individual differences. This decomposition provides a principled foundation for subsequent analyses of the spatial organization of cortical oscillatory modes and their relationships with multimodal brain measures.

#### fMRI data preprocessing and functional gradient estimation

Raw functional MRI data were converted from DICOM format into the Brain Imaging Data Structure (BIDS)(*22*) standard using HeuDiConv (v0.13.1) (*23*). Structural and functional preprocessing was subsequently carried out with fMRIPrep (v23.0.2) (*24*), implemented within the Nipype (v1.8) workflow engine (*25*). Anatomical preprocessing comprised intensity non-uniformity correction, skull removal, tissue segmentation, cortical surface reconstruction, and normalization to MNI space. Functional preprocessing included head-motion correction, slice timing adjustment, and alignment to the corresponding T1-weighted anatomical image.

Preprocessed functional time series were parcellated according to the Schaefer 200-region atlas (*26*) with seven-network resolution. Denoising was performed using the “simple” confound regression strategy, incorporating high-pass temporal filtering, removal of motion-and tissue-related nuisance signals, linear detrending, and z-score normalization (*27*).

Regional functional connectivity matrices were constructed for each participant by calculating zero-lag Pearson correlation coefficients between mean regional time courses.

Functional connectivity gradients were derived using the BrainSpace toolbox, applying the same parameter settings as those used for MEG-based gradient estimation. Prior to gradient computation, connectivity matrices were z-transformed and sparsified by retaining the strongest 10% of connections for each region, following established best practices. A cosine similarity kernel was used to construct an affinity matrix capturing similarities between regional connectivity profiles. Diffusion map embedding (*28*) was then employed to obtain low-dimensional representations of individual connectomes. The first ten gradients were extracted, and the principal gradient—corresponding to the canonical sensory-to-association axis—was selected for downstream analyses. To ensure comparability across participants, individual gradients were aligned to a group-level template derived from 100 unrelated Human Connectome Project participants using Procrustes alignment (*29*).

#### Diffusion MRI preprocessing and structural gradient estimation

Diffusion-weighted imaging data from the Cam-CAN cohort were processed using QSIPrep (v0.17.0) (*30*). The preprocessing pipeline included thermal noise reduction, correction for Gibbs ringing, bias field correction, intensity normalization, motion and eddy current correction, and spatial normalization to standard space.

Structural connectomes were reconstructed using the *mrtrix_multishell_msmt_ACT-hsvs* workflow. Fiber orientation distributions were estimated via multi-shell multi-tissue constrained spherical deconvolution (MSMT-CSD) (*31*). Whole-brain probabilistic tractography was performed with the iFOD2 algorithm under anatomically constrained tractography (ACT), informed by T1-weighted tissue segmentation. Streamline weights were refined using the SIFT2 algorithm, yielding weighted structural connectivity matrices (*32*).

Structural connectivity gradients were computed using the BrainSpace toolbox with default parameters. Structural connectivity matrices were first z-scored and thresholded to retain the top 10% of strongest connections per region. Cosine similarity was used to generate an affinity matrix representing similarity between regional connectivity profiles. Diffusion map embedding was subsequently applied to extract low-dimensional manifold representations of individual structural connectomes. The first three gradient components were retained for subsequent analyses. To align gradients across individuals, Procrustes transformation was applied using a reference template generated from the group-averaged structural connectivity matrix.

#### Structural MRI preprocessing and geometric gradient estimation

High-resolution T1-weighted anatomical images were processed using FreeSurfer’s standard *recon-all* pipeline, including skull stripping, intensity normalization, surface reconstruction, and cortical parcellation. Individual white matter surfaces were extracted as triangulated meshes, resampled to approximately 164,000 vertices, and registered to the *fsaverage* template through spherical alignment. Visual inspection and quality control procedures were conducted to ensure topological accuracy. Geometric gradients were derived by computing eigenmodes of the Laplace–Beltrami operator (LBO) on the cortical surface (*33*). The LBO was discretized on the *fsaverage* mesh using a cotangent-weighted formulation applied to triangular faces. Eigenmodes were obtained by solving the Helmholtz equation, Δψ = −λψ, under Neumann boundary conditions, where Δ denotes the discrete LBO, λ the eigenvalues, and ψ the corresponding eigenfunctions. These eigenfunctions form an ordered basis of spatially smooth patterns on the cortical manifold, ranging from low- to high-frequency spatial variations. The lowest non-trivial eigenmode corresponds to the smoothest spatial gradient across the cortex, whereas higher-order modes encode progressively finer spatial detail. In this study, the first 200 non-constant eigenmodes were retained, providing a multiscale geometric basis for projecting and analyzing cortical maps. This approach offers anatomically grounded, orthogonal, and scale-resolved representations of cortical organization.

#### Wilson–Cowan modeling of large-scale brain dynamics

The Wilson–Cowan (WC) model provides a canonical description of population-level neural dynamics through interactions between excitatory and inhibitory neuronal populations (*34*). The model incorporates local excitatory–inhibitory coupling, delayed network interactions governed by structural connectivity and conduction delays, nonlinear population responses implemented via sigmoidal transfer functions, refractory dynamics, and time constants controlling temporal evolution.

Empirical features were extracted from MEG recordings by parameterizing regional power spectra and estimating center frequency for each brain region (*35*). Center frequency was selected as the target feature due to its interpretability and robustness, reflecting both local excitation–inhibition balance and large-scale network effects. The WC model was embedded within individualized structural connectivity matrices to constrain network interactions (*36*).

Model parameters, including global coupling strength and signal propagation velocity, were optimized to reproduce the spatial distribution of empirical center frequencies. Model performance was quantified as the correlation between simulated and empirical regional center-frequency vectors. Accordingly, optimization focused on matching the spatial pattern of center frequencies rather than their absolute magnitudes. A detailed description of the parameter search and optimization procedure is provided in the *Supplementary Materials* (“Parameter Setting for Wilson–Cowan Modeling”).

#### Cortical annotations

To physiologically contextualize the common cortical oscillatory modes, we examined their relationships with normative maps of cytoarchitecture, cellular composition, and neurotransmitter systems. Cytoarchitectural features were quantified using BigBrain intensity profiles generated with the BigBrainWarp toolbox (*37*, *38*). These profiles captured variations in cell-body staining intensity sampled across 50 equivolumetric cortical surfaces, enabling laminar-sensitive characterization of cortical microstructure (*39*, *40*).

Common cortical oscillatory modes were correlated with spatial maps representing cellular composition derived from prior work (*41*). Molecular signatures for distinct cell classes were obtained from single-nucleus Drop-seq (snDrop-seq) datasets (*42*), and regional cell-type proportions were estimated via transcriptomic deconvolution of microarray data from the Allen Human Brain Atlas (AHBA) (*43*). The resulting dataset comprised 24 cell classes, encompassing excitatory and inhibitory neuronal subtypes as well as glial and vascular populations.

Neurotransmitter receptor and transporter distributions were assessed using positron emission tomography (PET) tracer maps covering 18 molecular targets across nine major neurotransmitter systems, curated by (*44*, *45*) (https://github.com/netneurolab/hansen_receptors). These systems included dopaminergic, noradrenergic, serotonergic, cholinergic, glutamatergic, GABAergic, histaminergic, cannabinoid, and opioid pathways, consistent with established neurochemical mapping frameworks (*46*, *47*). PET images were nonlinearly registered to the MNI-ICBM 152 (2009c asymmetric) template and subsequently parcellated using the Schaefer 200-region atlas. For molecular targets represented by multiple PET tracers, weighted averaging was employed to derive a single composite map per receptor or transporter.

#### Null model

To assess the spatial correspondence between common oscillatory modes and other neurobiological features, we employed a spatial permutation–based null model that accounts for spatial autocorrelation inherent to cortical maps. Neurobiological feature maps were first aligned to the oscillatory statistical maps, and their spatial correlations were computed.

Null distributions were generated by applying spherical rotations to cortical surface maps, thereby preserving local spatial continuity while disrupting the original spatial alignment between maps. After rotation, parcel values were reassigned based on the nearest rotated cortical region. This procedure was repeated 1,000 times to obtain a null distribution of correlation values (*46*). Rotations were performed on one hemisphere and mirrored to the contralateral hemisphere to maintain interhemispheric symmetry.

Statistical significance was determined by comparing the empirical correlation against the null distribution, with associations deemed significant if they exceeded the *95th* percentile of correlations derived from both spatial and temporal permutation procedures.

#### Age effect

Furthermore, we leveraged the Cam-CAN dataset to examine lifespan-wide age-related variations in spectral characteristics. Specifically, we assessed the association between age and center frequency as well as loading strength using partial Pearson correlations, controlling for handedness and sex, and applied false discovery rate (FDR) correction for multiple comparisons.

#### Test–retest reliability assessment analysis

To assess the test–retest reliability of the neurophysiological gradients, we quantified reliability using the intraclass correlation coefficient (ICC). By definition, ICC captures the relative contributions of within-subject and between-subject variability, such that higher reliability reflects lower within-subject variability and/or greater between-subject variability. Because reliability places an upper bound on the validity of metrics used for clinical inference, it constitutes a fundamental prerequisite for biomarker development in applied and translational contexts. In addition, we reconstructed the frequency components corresponding to the common oscillatory modes derived from the Cam-CAN dataset within the OMEGA dataset and quantified their test–retest reliability by computing intraclass correlation coefficients (ICCs) across multiple recording sessions.

#### Applications in Parkinson’s Disease

We used the CAM-CAN database to derive subject-specific shared subspaces and to reconstruct frequency loadings for individuals with Parkinson’s disease (PD) and HC in the OMEGA cohort. We then quantified the center frequency and loading strength. Finally, we performed two-sample *t*-tests to compare group differences in changes in center frequency or loading strength between patients and controls, with handedness, sex, and age included as covariates.

## DATA AVAILABILITY

MEG, fMRI, DWI data could be available on https://cam-can.mrc-cbu.cam.ac.uk/dataset/. OMEGA dataset could be available on https://www.mcgill.ca/bic/neuroinformatics/omega. Public resources used in this study can be accessed online, including the Allen Human Brain Atlas (AHBA; http://human.brain-map.org/), neuromaps (https://netneurolab.github.io/neuromaps/usage.html), BrainSpace (https://brainspace.readthedocs.io/en/latest/), and the ENIGMA Toolbox (https://enigma-toolbox.readthedocs.io/en/latest/pages.html).

## CODE AVAILABILITY

Codes will be available upon reasonable request.

## ACKNOWLEDGMENTS

Xiaobo Liu is supported by the China Scholarship Council.

## Wilson–Cowan Model Parameterization and Simulation

In the structural connectivity and preprocessing stage, we employed an individual-level structural connectome based on the Schaefer-200 cortical parcellation. The fiber-count matrix was max-normalized and used as the interregional coupling (weight) matrix, while the mean fiber-length matrix was used to estimate interregional distances. All primary analyses were conducted on individual connectomes rather than group-averaged networks.

Local neural dynamics within each brain region were modeled using a Wilson–Cowan two-population framework, explicitly representing interacting excitatory and inhibitory neuronal populations. Each node incorporated two classes of local coupling terms: within-population (self–self) interactions and cross-population (self–other) interactions. Long-range interregional communication was implemented through time-delayed network interactions, in which inputs from other regions were weighted by the structural connectivity matrix and propagated with transmission delays determined by fiber lengths and a fixed conduction velocity. Both excitatory and inhibitory populations received these long-range inputs.

Model parameters, search space, and external drives were defined as follows. The excitatory local gain was treated as a free parameter and sampled within the range 0.8–1.2, while the inhibitory gain was fixed at one quarter of the excitatory gain. The delay-related time constant was varied between 12 and 14, and the global coupling strength (network gain) was also sampled within the range 12–14. External drives to the excitatory and inhibitory populations were specified by the four-tuple [1,0.75,1,T][1, 0.75, 1, T][1,0.75,1,T], where 0.75 denotes an amplitude scaling factor and TTT corresponds to the total simulation duration. This external input was applied identically to all 200 nodes. Parameter estimation was performed using a uniformly sampled random grid across the defined parameter ranges.

For numerical simulations, each parameter set was applied to the whole-brain network to generate excitatory population time series for all nodes. The system was integrated with a time step of 0.5 ms for a total of 30,000 steps. The sampling rate followed the native implementation of the model, and no additional resampling was performed. Power spectral density was computed for the excitatory time series of each node, and the peak frequency within the 4–30 Hz band was identified as the node-specific center frequency. These values were assembled into a simulated center-frequency vector, which was compared against an empirical center-frequency vector derived from MEG data.

For model fitting and optimization, the objective function targeted the spatial distribution of center frequencies rather than time-domain signal similarity. Specifically, the Pearson correlation coefficient between the empirical and simulated center-frequency vectors was used as the fitting metric. This approach emphasizes spatial concordance and robustness across regions, rather than minimizing per-node amplitude discrepancies. A random grid search consisting of 500 parameter samples was performed. For each sample, simulations and feature extraction were carried out, followed by computation of the correlation coefficient.

The parameter set yielding the highest correlation was selected as the optimal solution, and both the optimal parameters and the corresponding goodness-of-fit value were recorded.

**Supplementary Fig. 1.**
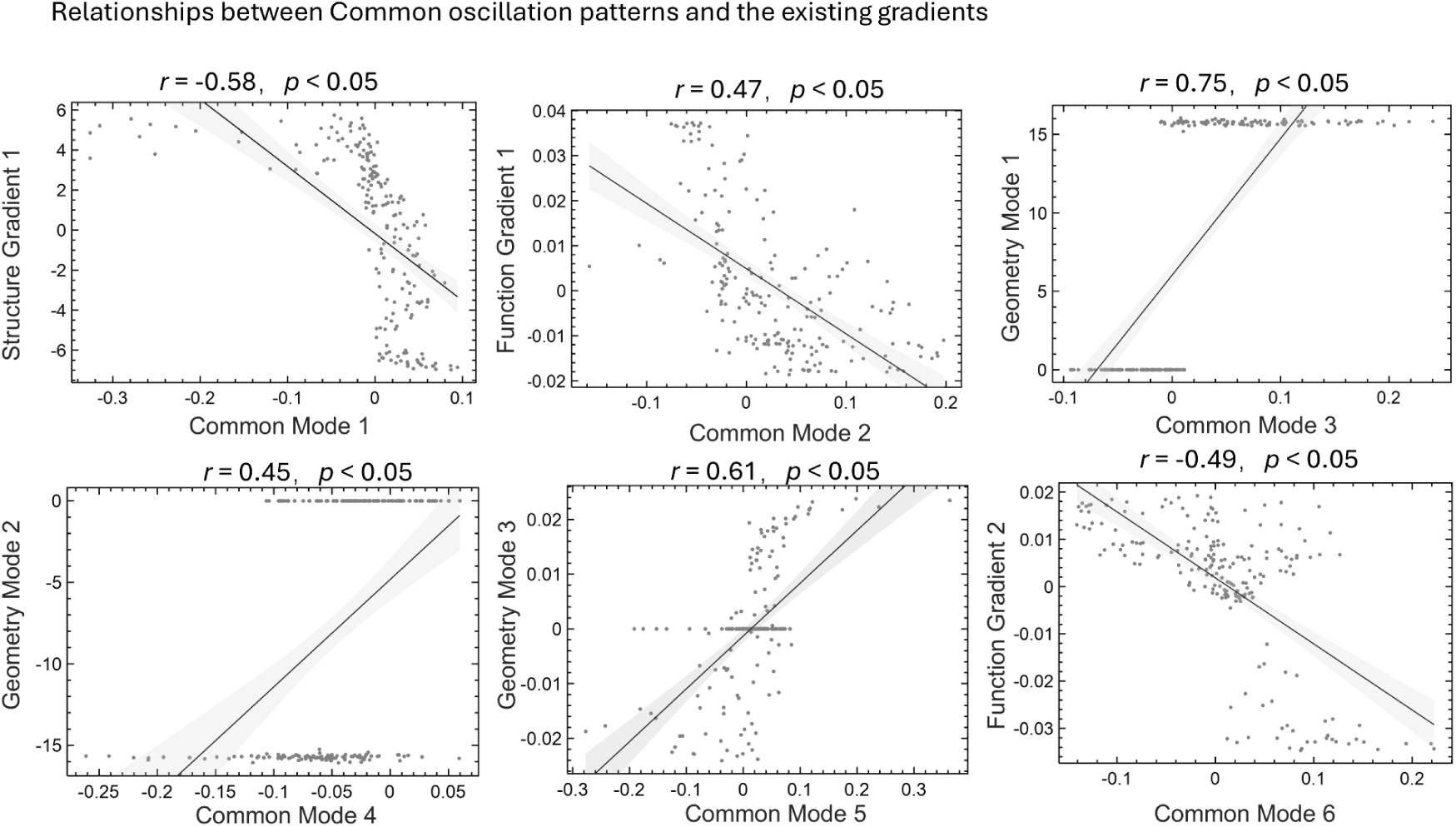
Statistical relationships between common oscillatory modes and established brain gradients. Scatter plots illustrate the spatial correlations between six common oscillatory modes and multiple brain gradients, including structural gradients, functional gradients, and geometric modes. Associations were quantified using Pearson’s correlation coefficient, with statistical significance assessed at p<0.05p < 0.05p<0.05. Solid black lines denote linear regression fits, and shaded areas indicate the 95% confidence intervals.

Common Mode 1 showed a significant negative correlation with Structure Gradient 1 (r=−0.58r = -0.58r=−0.58, p<0.05p < 0.05p<0.05). Common Mode 2 exhibited a significant positive correlation with Function Gradient 1 (r=0.47r = 0.47r=0.47, p<0.05p < 0.05p<0.05). Common Mode 3 was strongly and positively correlated with Geometry Mode 1 (r=0.75r = 0.75r=0.75, p<0.05p < 0.05p<0.05). Common Mode 4 demonstrated a significant positive association with Geometry Mode 2 (r=0.45r = 0.45r=0.45, p<0.05p < 0.05p<0.05). Common Mode 5 showed a significant positive correlation with Geometry Mode 3 (r=0.61r = 0.61r=0.61, p<0.05p < 0.05p<0.05). Common Mode 6 was significantly negatively correlated with Function Gradient 2 (r=−0.49r = -0.49r=−0.49, p<0.05p < 0.05p<0.05).

Collectively, these results indicate that distinct whole-brain oscillatory patterns are spatially aligned with structural, functional, and geometric principles of brain organization, highlighting a multidimensional coupling between neural dynamics and macroscale cortical architecture.

